# 3D-Manhattan: An interactive visualization tool for multiple GWAS results

**DOI:** 10.64898/2026.03.14.709978

**Authors:** Shumpei Hashimoto

## Abstract

Genome-wide association studies (GWAS) are widely used to identify genetic loci underlying various agronomic traits. Conventional Manhattan plots provide an effective two-dimensional (2D) summary of an individual GWAS result. However, recent advances in high-throughput phenotyping have led to study designs that generate multiple GWAS outputs across time points, traits, or experimental conditions. In such settings, biological insight increasingly depends on comparative interpretation of multiple association maps, yet panel-based arrangements of 2D plots fragment related information and impede recognition of shared or dynamic genetic signals. Here, I present 3D-Manhattan, an interactive visualization framework that integrates multiple GWAS results within a unified three-dimensional (3D) coordinate system. By extending the conventional Manhattan plot with an additional axis representing time, trait, or condition, 3D-Manhattan enables simultaneous, axis-aligned comparison of association landscapes while preserving genomic coordinates and statistical values. The tool is implemented as a stand-alone, browser-based application using WebGL-based rendering and supports smooth interaction without server-side computation. The framework provides flexible visualization controls, region highlighting, and variant-level correspondence across datasets, facilitating exploratory analysis of stable and context-dependent genetic associations. Collectively, 3D-Manhattan provides an alternative approach for visualizing multi-dimensional GWAS results and offers a powerful platform for visualizing general a series of genome-wide datasets.

## Introduction

Genome-wide association studies (GWAS) have become a central analytical framework for dissecting the genetic basis of quantitative traits across a wide range of biological systems. By testing statistical associations between genome-wide genetic variation and phenotypic diversity, GWAS have enabled the identification of loci underlying agronomic performance, physiological processes, and developmental regulation (Hamblin et al., 2011; Huang and Han, 2014; Lipka et al., 2015; Myles et al., 2009). The widespread availability of high-density genotyping platforms and reference genomes has further established GWAS as a routine and scalable approach in both model organisms and crop species (Huang et al., 2013; Nguyen et al., 2019). Traditionally, the visualization of GWAS results has relied on Manhattan plots, which provide a compact two-dimensional (2D) representation of association signals along genomic coordinates (D. Turner, 2018; LiLin-Yin, 2024). This visualization paradigm has proven highly effective for highlighting genomic regions that exceed statistical significance thresholds and for facilitating rapid candidate locus identification. As a result, Manhattan plots have become a de facto standard for reporting GWAS results and are supported by a broad ecosystem of software tools.

However, the landscape of GWAS applications has changed substantially in recent years. Advances in high-throughput phenotyping have dramatically expanded both the temporal resolution and dimensionality of phenotypic data (Yang et al., 2020). Imaging-based phenotyping, continuous sensor measurements, and automated experimental platforms now routinely generate dense time-series data and multi-trait measurements across developmental stages or environmental conditions (Bac-Molenaar et al., 2015; Camargo et al., 2018; Neumann et al., 2017, 2015; Pauli et al., 2016; Ren et al., 2018; Wang et al., 2015). These advances have given rise to time-resolved and multi-dimensional GWAS designs, in which association analyses are performed repeatedly across time points, traits, or experimental contexts. In such studies, the analytical challenge is no longer limited to interpreting a single association map. Instead, biological insight increasingly depends on the comparative interpretation of multiple GWAS outputs: identifying loci whose effects emerge, disappear, or shift over time; distinguishing stable genetic signals from context-dependent associations; and recognizing global patterns that span multiple analyses. In practice, these results are often visualized as collections of separate Manhattan plots arranged across panels or figures. While this approach preserves the integrity of individual analyses, it fragments related information across disconnected visual spaces and places a growing cognitive burden on the viewer as the number of plots increases. From a visualization perspective, this fragmentation represents a fundamental limitation of 2D representations when applied to inherently multi-dimensional data. When association results are distributed across time, traits, or experimental conditions, relationships among datasets are not explicitly encoded within a shared coordinate system. Therefore, subtle but biologically meaningful patterns—such as gradual shifts in association strength or coordinated changes across loci—may be difficult to recognize through sequential inspection alone.

Here, I present 3D-Manhattan, an interactive visualization framework designed to address this limitation by embedding multiple GWAS results within a three-dimensional (3D) space. By extending the conventional Manhattan plot with an additional axis representing time, trait, or experimental condition, 3D-Manhattan enables multiple association landscapes to be examined simultaneously within a unified coordinate system. This representation preserves genomic context while enhancing the visual continuity across datasets. Coupled with an interactive user interface that supports dynamic manipulation of viewing parameters, 3D-Manhattan facilitates exploratory analysis of complex GWAS results and provides an intuitive means to uncover patterns that are difficult to discern using conventional 2D visualizations.

## Materials and Methods

### Implementation of 3D-Manhattan

3D-Manhattan was developed on an Apple Mac Studio with an Apple M4 Max processor and 36 GB RAM. The latest version of 3D-Manhattan is available on the GitHub repository (https://github.com/hashimotoshumpei/3D-Manhattan). The repository includes detailed instructions on how to use the tool. An interactive AI-assisted coding workflow was partially utilized during development to facilitate rapid prototyping and iterative UI refinement, with all code reviewed and finalized by the author. 3D-Manhattan is implemented as a browser-based stand-alone visualization tool using JavaScript and WebGL (via the Three.js library).

## Results

### Concept and Implementation Overview

This tool is designed to integrate multiple genome-wide association study (GWAS) results derived from serial or multi-dimensional datasets, including time-course phenotypes, multiple traits, or model evaluation outputs (Fig.1). In this workflow, each dataset is independently subjected to GWAS, generating standard association result files that are subsequently imported into the 3D-Manhattan interface (Fig.1). These datasets are then processed and rendered within a unified 3D coordinate system, in which individual Manhattan plots are stacked along an additional axis representing time points, traits, or experimental conditions. Importantly, genomic coordinates and −log_10_(p) values are preserved across all datasets, enabling direct, axis-aligned comparison within a consistent genomic reference frame. The interactive interface allows users to dynamically control key visualization parameters, such as chromosome selection, inter-dataset spacing, point size, and axis scaling, facilitating both global overview and detailed inspection of association patterns. As illustrated in Fig. 1, the framework supports multiple output modes, including genome-wide stacked Manhattan plots, chromosome-specific visualizations, and variant-level connections across datasets. Thus, these features enable simultaneous exploration of multiple GWAS results within a single spatial context, providing a structured and intuitive representation of complex, multi-dimensional association data.

**Fig. 1.**
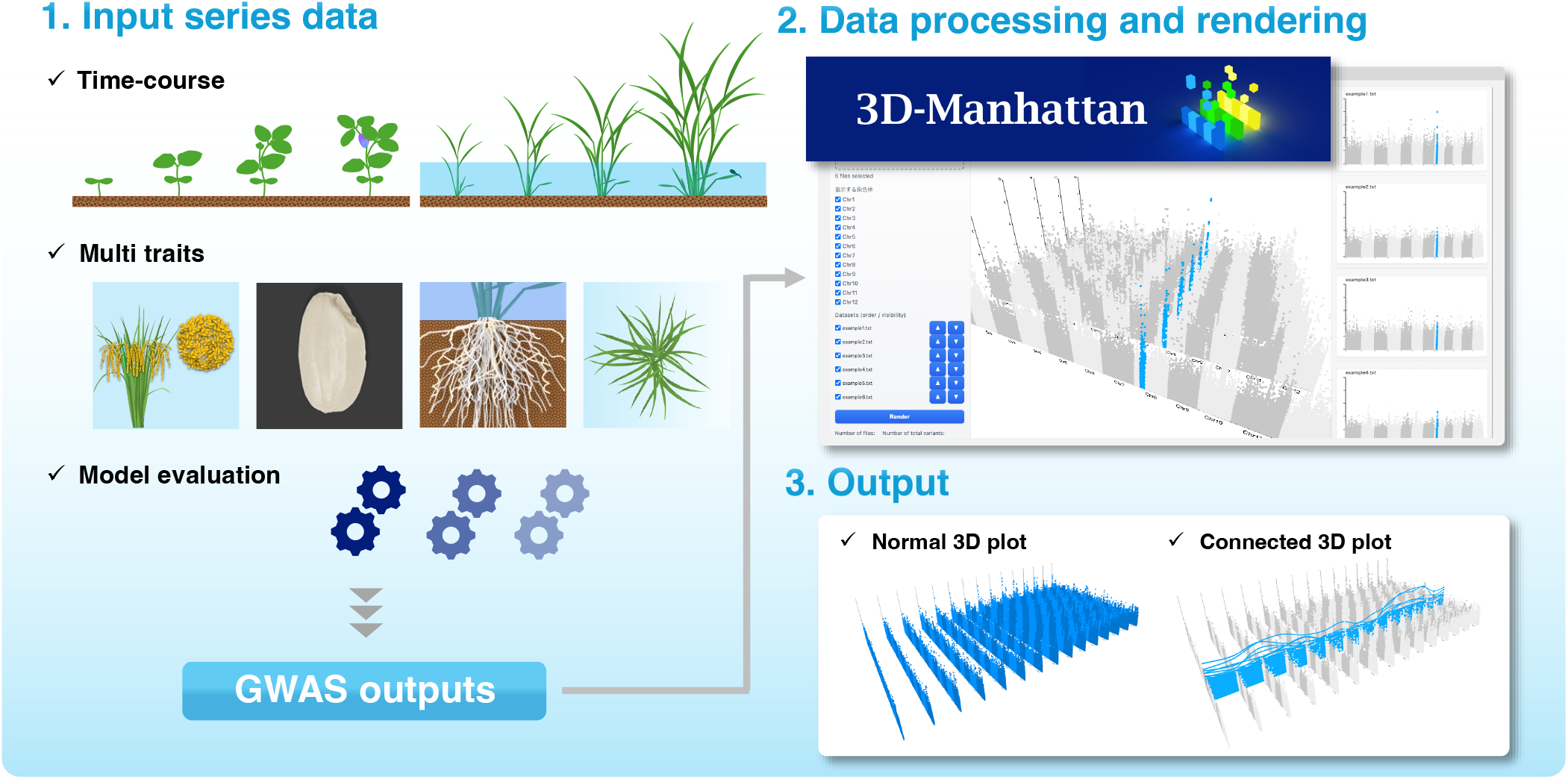
Overview of the concept and workflow of the 3D Manhattan visualization tool. In the first step, time-series phenotypic data, multiple traits, or model evaluation results are used as inputs to genome-wide association studies (GWAS). GWAS outputs from multiple datasets are processed and rendered using the 3D Manhattan visualization interface in the second step, which allows interactive control of parameters such as chromosome selection, stack spacing, point size, and axis scaling. Finally, the tool provides flexible output views, including genome-wide stacked Manhattan plots and variant-to-variant connections across datasets, enabling intuitive exploration of complex multi-dimensional GWAS results.

### User Interface (UI) and User Experience (UX)

The tool operates locally within modern web browsers and does not require server-side computation or installation of external dependencies. GWAS results are supplied as tab-separated value (TSV) files containing chromosome identifiers, base-pair positions, and association statistics (e.g., −log_10_(p) values). For each dataset, chromosomes are mapped to continuous genomic coordinates according to their physical lengths, consistent with conventional Manhattan plot representations, thereby enabling genome-wide visualization with proportional spacing. A distinctive feature of this tool is that multiple GWAS datasets are stacked along the z-axis to generate a 3D Manhattan plot. The 3D view supports interactive manipulation, allowing users to adjust the viewing angle via mouse dragging and to control zoom levels using the mouse wheel. Visualization parameters, including vertical scaling, point size, inter-dataset spacing, axis offsets, and significance threshold lines, can be interactively adjusted through a graphical UI. Chromosome colors can also be customized, with independent color assignment for odd- and even-numbered chromosomes. The 3D plot, rendered at an arbitrary viewpoint, can be exported at high resolution in PNG, JPEG, or PDF formats from any selected viewpoint. Furthermore, conventional 2D Manhattan plots corresponding to individual datasets are displayed alongside the 3D visualization, enabling detailed inspection while preserving a genome-wide overview (Fig. 2). Together, these features provide a flexible and intuitive UI that accommodates varying dataset sizes and diverse analytical objectives.

**Fig. 2.**
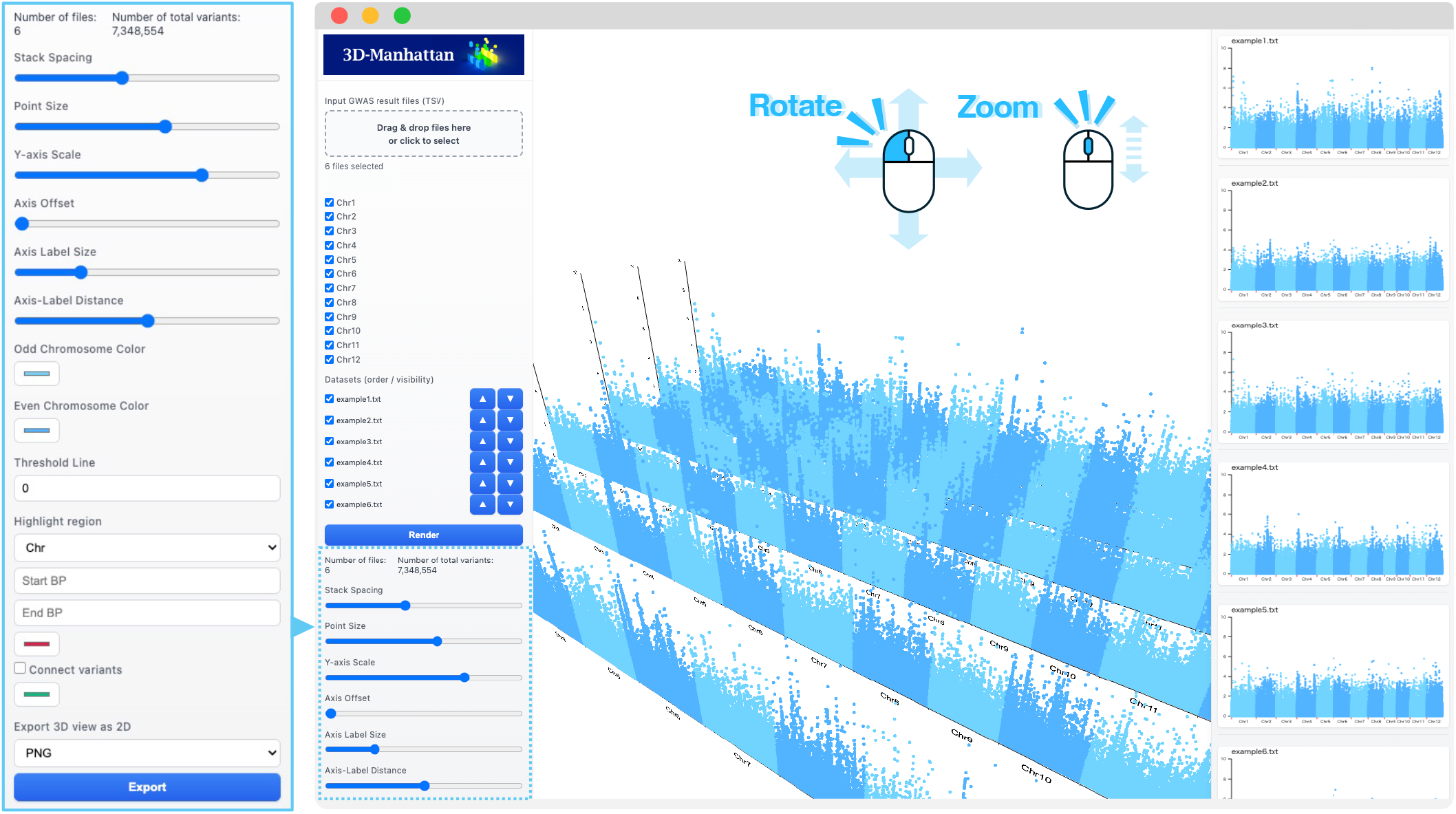
Graphical user interface and interactive visualization of multi-dataset GWAS results. Multiple GWAS result files can be loaded simultaneously, and users can interactively select chromosomes and adjust visualization parameters such as stack spacing, point size, y-axis scale, axis offset, and label size. The central panel displays stacked 3D Manhattan plots representing multiple datasets, while the right panel provides corresponding 2D Manhattan plots for each dataset, allowing simultaneous overview and detailed inspection of GWAS signals.

### Genome-wide and chromosome-level comparison

A genome-wide view of stacked GWAS results is shown in Fig. 3A, enabling direct comparison of association signals across multiple datasets. Shared association peaks across datasets are readily identified as vertically aligned signals, whereas dataset-specific peaks appear as isolated features. The tool also supports focused visualization of individual chromosomes as well as alternative chromosome arrangements, facilitating detailed examination of regions of interest (Fig. 3B). Such views are particularly useful for exploring temporal changes in association strength or trait-specific genetic effects across datasets.

**Fig. 3.**
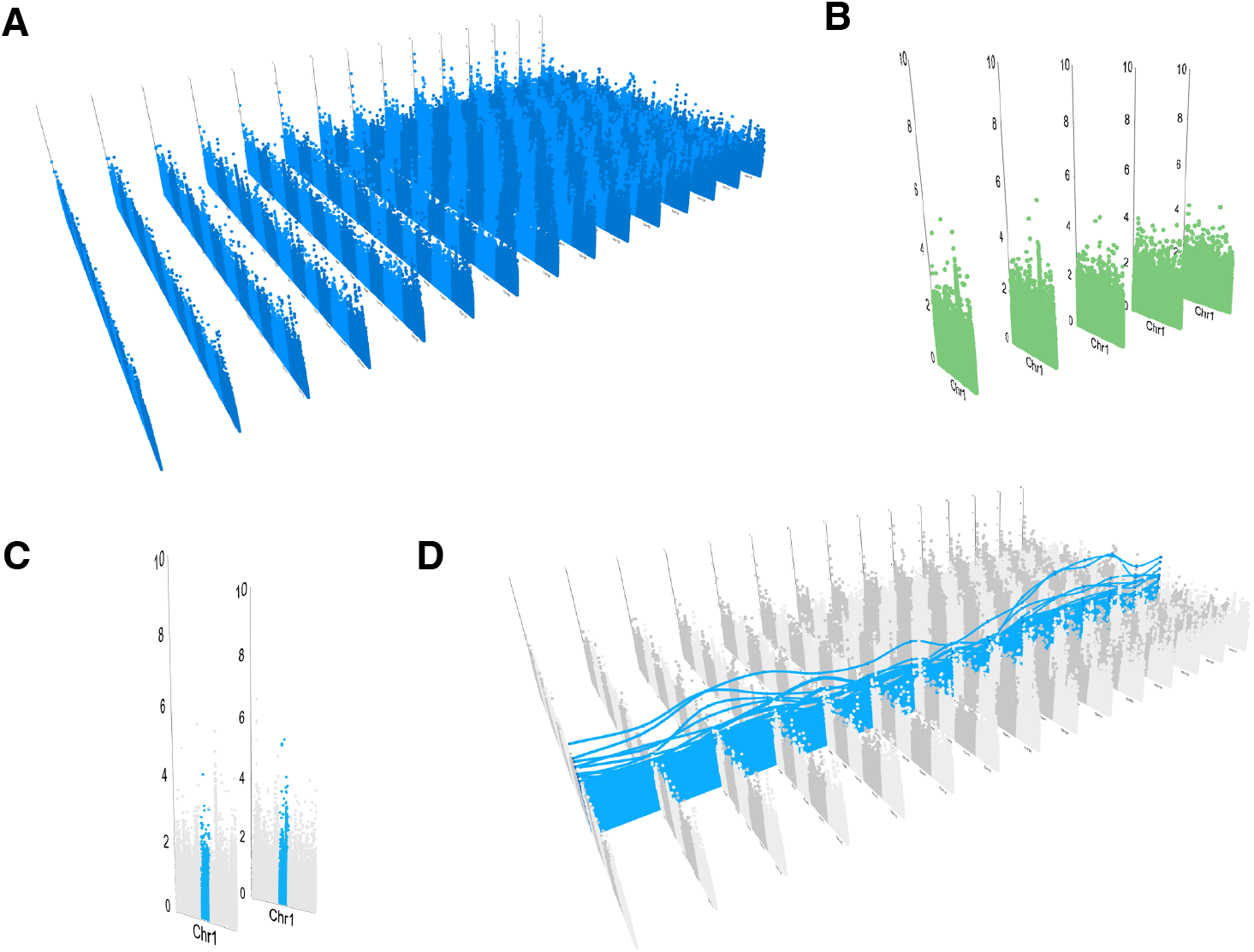
Example visualization generated by 3D-Manhattan. (A) Genome-wide visualization showing multiple datasets stacked along the depth axis, enabling comparison of association patterns across all chromosomes simultaneously. (B) Zoomed view of a selected chromosome across multiple datasets, highlighting differences in signal intensity and distribution. (C) 3D Manhattan plots for two datasets with highlighted genomic regions. (D) Variant-to-variant connections between datasets are highlighted using linking lines, enabling direct comparison of corresponding variant signals across datasets for the same chromosome.

### Visualization of variant correspondence across datasets

3D-Manhattan further provides a highlighting function for specific genomic regions or individual variants. By selecting a chromosome and specifying genomic coordinates, users can assign an arbitrary color to the corresponding region (Fig. 3C). To highlight a single polymorphic site, the same genomic position is specified as both the start and end coordinates. Multiple regions can be highlighted simultaneously, and highlighted regions are reflected in real time in both the three-dimensional and two-dimensional plots. A distinctive feature of 3D-Manhattan is its ability to visually link variants located in similar genomic regions across different datasets (Fig. 3D). By drawing connecting lines between corresponding regions, the tool emphasizes reproducible signals and facilitates identification of stable or dynamic genetic associations across conditions. The color of the connecting lines can be freely customized by the user. This functionality is particularly valuable for visualizing serial GWAS results, in which the persistence or disappearance of association signals across developmental stages provides insight into stage-specific genetic regulation.

## Discussion

The visualization of multiple GWAS results has been commonly achieved by arranging 2D Manhattan plots across panels or stacking them vertically. While effective for presenting individual analyses, this approach fragments related datasets into separate visual contexts. Consequently, comparisons across time points, traits, or experimental conditions rely on sequential inspection. Continuity across datasets is therefore not explicitly encoded within a unified coordinate system. As the number of GWAS outputs increases, such panel-based representations impose a growing cognitive burden and limit efficient comparative exploration.

3D visualization offers a principled solution to this limitation by enabling multiple analytical results to be represented within a shared spatial framework. In the context of GWAS, introducing a third axis corresponding to time, trait, or experimental condition allows multiple Manhattan plots to be embedded within a single coordinate space. This integration preserves genomic position and signal intensity. Furthermore, it enables users to examine relationships among datasets simultaneously. As a result, global patterns, gradual transitions, and condition-specific dynamics can be detected more readily than with isolated two-dimensional views.

Several tools have explored the use of 3D representations for GWAS data. PheGWAS, for example, visualizes genome-wide by phenome-wide association matrices as an interactive 3D landscape (George et al., 2020). This design enables users to explore associations across many phenotypes concurrently. It is particularly effective for identifying pleiotropic loci and cross-trait relationships. PheGWAS therefore illustrates the value of 3D visualization for summarizing large-scale association structures (George et al., 2020). Another notable tool, BigTop, represents a more immersive exploration paradigm by implementing GWAS visualization within a virtual reality environment (Westreich et al., 2020). In BigTop, Association signals are arranged around the user in a cylindrical space (Westreich et al., 2020). Chromosomal position is mapped circumferentially, −log10(p) is represented vertically, and an additional dimension such as minor allele frequency is encoded radially (Westreich et al., 2020). Through point-level interaction, users can inspect individual variants and their annotations. This design demonstrates how immersive depth perception and multidimensional encoding can enhance intuitive understanding of GWAS signal distributions.

Despite these advances, existing 3D tools are not primarily designed for the systematic comparison of multiple GWAS results generated under different conditions. In particular, neither phenome-wide overview landscapes nor immersive virtual environments explicitly preserve the conventional Manhattan plot structure required for direct, axis-aligned comparison across analyses. To address this gap, I developed 3D-Manhattan, an interactive visualization framework that integrates multiple GWAS outputs into a single 3D coordinate space (Fig.1). Importantly, this integration is achieved while retaining the familiar Manhattan plot representation. By stacking individual Manhattan plots along a third axis representing time points, traits, or experimental conditions, 3D-Manhattan enables direct visual tracking of how association signals emerge, shift, or disappear across analyses (Fig.2). This tracking is performed within a consistent genomic reference frame, facilitating intuitive comparison across conditions (Fig.3).

The technical foundation of 3D-Manhattan is its implementation using three.js, a JavaScript framework built on Web Graphics Library (WebGL) for interactive 3D graphics (Cabello, R. and three. js Contributors, 2010). Originally released in 2010, three.js was developed to simplify WebGL-based rendering and has since become widely adopted for interactive visualization in modern web browsers, including WebXR-based virtual and augmented reality applications. In contrast, widely used GWAS visualization tools and plotting libraries, such as qqman (D. Turner, 2018), CMplot (LiLin-Yin, 2024), matplotlib (Hunter, 2007), and ggplot2 (Wickham, 2016), are primarily optimized for static, CPU-based 2D visualization. Therefore, they scale poorly to interactive 3D exploration of genome-wide datasets. By leveraging GPU-accelerated rendering through WebGL, three.js enables efficient real-time interaction directly within web browsers. As a result, 3D-Manhattan maintains smooth and responsive performance even when handling large-scale GWAS datasets.

By combining an axis-preserving 3D design with a stand-alone, browser-based implementation, 3D-Manhattan provides a complementary visualization paradigm for multi-condition and multi-trait GWAS results. Notably, although the visualization framework presented here was developed for 3D Manhattan plots, it naturally suggests broader applicability to the visualization of other forms of serial biological data.

## Author Contribution Statement

SH developed the software and wrote the manuscript.

## Acknowledgments

I express my gratitude to Mr. Kentaro Yamada, Dr. Takeshi Izawa, Dr. Ammar Elakhdar, and Mr. Takumi Kawauchi from the Laboratory of Plant Breeding and Genetics, Department of Agricultural and Environmental Biology, University of Tokyo, for their valuable contributions to this work. Mr. Yamada provided insightful code reviews and suggestions, Dr. Izawa engaged in in-depth discussions. Dr. Ammar Elakhdar and Mr. Takumi Kawauchi provided valuable comments on the manuscript. This research was partially supported by JSPS KAKENHI Grant Number JP23KJ0326.

## Notes

### Competing Interest Statement

The authors have declared no competing interest.

